# A remarkably specific ligand reveals ghrelin *O*-acyltransferase interacts with extracellular peptides and exhibits unexpected cellular localization for a secretory pathway enzyme

**DOI:** 10.1101/2021.06.01.446150

**Authors:** Maria B. Campaña, Tasha R. Davis, Elizabeth R. Cleverdon, Michael Bates, Nikhila Krishnan, Erin R. Curtis, Marina D. Childs, Mariah R. Pierce, Sadie X. Novak, Yasandra Morales-Rodriguez, Michelle A. Sieburg, Heidi Hehnly, Leonard G. Luyt, James L. Hougland

**Affiliations:** Department of Chemistry, Syracuse University, Syracuse, NY 13244, USA; Department of Biology, Syracuse University, Syracuse, NY 13244, USA; Department of Chemistry, University of Western Ontario, London, Ontario, Canada N6A 2K7; BioInspired Syracuse, Syracuse University, Syracuse, NY 13244; Department of Oncology, and Department of Medical Imaging, London Regional Cancer Program, Lawson Health Research Institute, 800 Commissioners Road East, London, Ontario, Canada N6A 5W9

**Keywords:** ghrelin, ghrelin O-acyltransferase, membrane-bound O-acyltransferase, substrate inhibitor, GHS-R1a, acyltransferase, post-translational modification (PTM), membrane enzyme, prostate cancer, protein acylation

## Abstract

Ghrelin *O-*acyltransferase (GOAT) plays a central role in the maturation and activation of the peptide hormone ghrelin, which performs a wide range of endocrinological signaling roles. Using a tight-binding fluorescent ghrelin-derived peptide designed for high selectivity for GOAT over the ghrelin receptor GHS-R1a, we demonstrate that GOAT interacts with extracellular ghrelin and facilitates ligand cell internalization in both transfected cells and prostate cancer cells endogenously expressing GOAT. Coupled with enzyme mutagenesis, ligand uptake studies provide the first direct evidence supporting interaction of the putative histidine general base within GOAT with the ghrelin peptide acylation site. Our work provides a new understanding of GOAT’s catalytic mechanism, establishes a key step required for autocrine/paracrine ghrelin signaling involving local reacylation by GOAT, and raises the possibility that other peptide hormones may exhibit similar complexity in their intercellular and organismal-level signaling pathways.

---

Ghrelin is a unique peptide hormone implicated in regulation of physiological pathways impacting appetite, energy storage and metabolism, neurological responses to stress, and reward processing associated with addictive behavior.^1–7^ Ghrelin exists in two distinct chemical forms in the bloodstream, acylated ghrelin with an eight-carbon fatty acid covalently linked to a serine side chain at position 3 and des-acyl ghrelin with a free serine hydroxyl at this position.^8^ Ghrelin is the only known peptide predicted to undergo this serine octanoylation modification, which is catalyzed by ghrelin *O*-acyltransferase (GOAT).^9^ Ghrelin octanoylation is required for binding to its cognate receptor the growth hormone secretagogue receptor type 1a (GHS-R1a), a member of the G-protein coupled receptor family.^7, 10^

Beyond their direct involvement in ghrelin maturation and signaling, the ghrelin binding proteins GHS-R1a and GOAT are potential disease biomarkers. The GHS-R1a receptor is being explored for cancer detection and imaging due to its altered expression in neoplasms including prostate, testicular, ovarian, breast, and neuroendocrine tumors.^11–13^ GOAT overexpression is also observed in multiple cancers including breast, endocrine tissue, and prostate.^14–18^ In a study accounting for metabolic alterations in prostate cancer (PCa) patients, GOAT was shown to be differentially expressed in these patients compared to non-cancer controls.^14^ GOAT has also been detected by ELISA-based assay in the urine and blood plasma of PCa patients, with GOAT plasma concentrations acting as a more consistent and sensitive detector of aggressive PCa than the traditional PSA diagnostic biomarker.^14–15^ To fully explore the potential for GOAT to serve as a novel disease biomarker, significant advancements are needed in the design of efficient probes for detecting GOAT and our understanding of this enzyme’s trafficking within both the cell and the body.

Originally annotated and observed in the endoplasmic reticulum membrane,^19^ GOAT has been recently suggested to also be distributed to the plasma membrane.^20–21^ Extracellular exposure on the plasma membrane would position GOAT to serve as a cancer cell biomarker through GOAT-specific ligands coupled to appropriate imaging groups. The high selectivity of GOAT for ghrelin, the only predicted GOAT substrate in the human proteome,^9^ supports the ability to design the potent GOAT-targeted ligands required to exploit this novel diagnostic and potential therapeutic target. In this work, we employed parallel structure-activity analyses to develop a synthetic ghrelin analog with exceptionally specific binding to GOAT over GHS-R1a. This ligand was further functionalized with a sulfo-Cy5 fluorophore to afford imaging of ligand binding to human cell lines expressing GOAT. The ligand probes designed in this study and our analysis of GOAT localization and ligand binding offer mechanistic insight into GOAT binding and catalytic strategies, demonstrates that GOAT interacts with extracellular peptides at the cell surface, and supports further exploration of GOAT as an imaging target for disease diagnostics.

## Results and Discussion

### Development of a specific high-affinity peptide ligand for GOAT

The high affinity and specificity of ghrelin binding to the GHS-R1a has enabled development of imaging agents targeting this receptor such as a fluoronaphthyl acylated ghrelin(1– 8) analogue for use in PET imaging.^22^ While GOAT can also bind ghrelin mimics acylated with fatty acids,^9, 23^ the affinity of these molecules for both GOAT and the GHS-R1a render these molecules unsuitable for specifically detecting and imaging GOAT in a cellular or organismal context. The GO-CoA-Tat bisubstrate GOAT inhibitor developed by Barnett and coworkers binds GOAT without exhibiting antagonism of the GHS-R1a receptor,^24^ demonstrating selectivity between these two ghrelin binding proteins. In GO-CoA-Tat, selectivity for GOAT is presumably due to inclusion of coenzyme A attached to the acyl side chain. To provide an easily functionalizable and synthetically accessible scaffold for ligand development, we explored a new class of GOAT ligands inspired by a class of substrate-mimetic GOAT inhibitors incorporating a free amino group in place of the serine hydroxyl at the acylation site.^25^ With these ligands lacking a hydrophobic moiety at the acylation site, we predicted they would exhibit significantly reduced binding affinity for the GHS-R1a.^10^ Compared to peptides acylated with either an octanoyl group (ligand **2**) or a 6-fluoro-2-naphthoyl group (ligand **3**), the peptide with a free Dap amino group (ligand **1**) exhibits tight binding to GOAT without detectible binding to the ghrelin receptor (Table 1 and Supporting Figure S1.).

**Table 1.** Ligand design and optimization for targeting the GHS-R1a and GOAT.

| **Ligand** | **Peptide Sequence** | **IC<sub>50</sub> against GHS-R1a** | **IC<sub>50</sub> against hGOAT** |
| --- | --- | --- | --- |
| *Acylation site modifications* |  |  |  |
| **1** | H-GS-Dap-FLSPY-NH<sub>2</sub> | >100 μM | 104 ± 43 nM |
| **2** | H-GS-(C8-Dap)-FLSPY-NH<sub>2</sub> | 65 nM<sup>a</sup> | 64 ± 21 nM |
| **3** | H-GS-(6FN-Dap)-FLSPY-NH<sub>2</sub> | 9.9 nM<sup>a</sup> | >100 μM |
| *Modifications at G1* |  |  |  |
| **4** | H-Aib-S-Dap-FLSPY-NH<sub>2</sub> | >100 μM | >100 μM |
| **5** | H-Inp-S-Dap-FLSPY-NH<sub>2</sub> | >100 μM | >100 μM |
| **6** | H-Aib-S-(6FN-Dap)-FLSPY-NH<sub>2</sub> | 9.2 nM<sup>a</sup> | ND |
| **7** | H-Inp-S-(6FN-Dap)-FLSPY-NH<sub>2</sub> | 9.6 nM<sup>a</sup> | ND |
| *Modifications at E8* |  |  |  |
| **8** | H-GS-(C8-Dap)-FLSPY-NH<sub>2</sub> | 65 nM<sup>a</sup> | 64 ± 21 nM |
| **9** | H-GS-(C8-Dap)-FLSPE-NH<sub>2</sub> | 200 nM<sup>a</sup> | 580 ± 60 nM |
| **10** | H-GS-(C8-Dap)-FLSPN-NH<sub>2</sub> | 31.9 nM<sup>a</sup> | 59 ± 7 nM |
| **11** | H-GS-(C8-Dap)-FLSPT-NH<sub>2</sub> | 3.3 nM<sup>a</sup> | 27 ± 8 nM |
| *Modifications at F4* |  |  |  |
| **12** | H-GS-Dap-Nal-1-LSPT-NH<sub>2</sub> | >100 μM | 6 ± 1 nM |
| **13** | H-GS-Dap-Nal-2-LSPT-NH<sub>2</sub> | >100 μM | 16 ± 5 nM |

<sup>a</sup> IC<sub>50</sub> values for ligands 2, 3, and 6-11 against GHS-R1a cited from Charron et al.<sup>22</sup>

Intriguingly, while the octanoylated ligand **2** binds tightly to both GOAT and the GHS-R1a, incorporation of the fluoronaphthoyl group at the acylation site in ligand **3** enhances receptor binding while blocking interaction with GOAT. This suggested the potential to tune ligand selectivity for both GOAT and the GHS-R1a to generate orthogonal ligands for each ghrelin-binding protein. Noting the strict selectivity for recognition of the *N*-terminal glycine residue (G1) by GOAT which requires both the *N*-terminal amino group and lack of side chain / steric bulk at the G1 position,^9, 25^ we replaced G1 with two unnatural amino acids - aminoisobutyric acid (Aib, ligand **4**) and isonipecotic acid (Inp, ligand **5**) - and determined binding affinities of these ligands for both GOAT and the GHS-R1a. Incorporation of either Aib or Inp at the peptide *N*-terminus had a pronounced negative impact on binding to GOAT while the GHS-R1a readily tolerated replacement of G1 in the context of acylated ligands **6** and **7**.^22^ Taken together, these studies identify the G1 position and the acylation site as the most important determinants for ligand specificity towards either GOAT or GHS-R1a (Table 1).

Having achieved ligand selectivity for GOAT over GHS-R1a through modifications at the acylation site, we sought to optimize ligand binding affinity for GOAT. We explored substitutions at glutamate 8 (E8) and phenylalanine 4 (F4) within the ghrelin-derived ligand based on previous studies demonstrating the involvement of these amino acids in ghrelin recognition by GOAT and GHS-R1a.^9, 22, 25–26^ Incorporation of a threonine residue at E8 strengthens ligand binding to the GHS-R1a, and we found this substitution similarly increases ligand binding to GOAT (ligand **11**) much more than either tyrosine or asparagine at this position. Both GOAT and GHS-R1a recognize the F4 residue,^22, 27^ with GOAT exhibiting a preference for large hydrophobic/aromatic amino acids at this position.^9, 25^ Exploiting this preference using unnatural amino acids, we found incorporation of 1-naphthylalanine (Nal-1, compound **12**) or 2-naphthylalanine (Nal-2, compound **13**) at the F4 position substantially increased ligand binding to GOAT in the context of unacylated ligands, which exhibited no detectible binding to the GHS-R1a (Table 1).

Correlating binding affinities for GOAT and GHS-R1a exhibited by ligands in this study highlights three classes of compounds, non-selective ligands and those with >100-fold selectivity for either GOAT or GHS-R1a (Figure 1). Combining the most successful substitutions in these studies has generated a highly selective ligand **12** for GOAT with nanomolar affinity, with ligand **14** previously reported as a highly potent ligand for GHS-R1a which combined elements of ligands **3** and **4** (Figure 2).^22^ Each of these ligands exhibits >15,000-fold selectivity for its target over the other ghrelin-binding protein. To equip the GOAT selective ligand for use in cell imaging, we synthesized ligand **15** containing a lysine residue at its *C*-terminus and a sulfo-Cy5 fluorophore attached to the lysine side chain (Figure 2). Earlier studies of labeled Cy5-ghrelin(1–19) demonstrated strong binding to the GHS-R1a, with less susceptibility to photobleaching and high detection of GHS-R expression in live differentiating cardiomyocytes.^28^ Ligand **15** exhibits potent binding to GOAT with a nanomolar IC_50_ value when assayed as an inhibitor, supporting our ability to functionalize GOAT peptide ligands with imaging groups without compromising GOAT binding ability.

**Figure 1.**
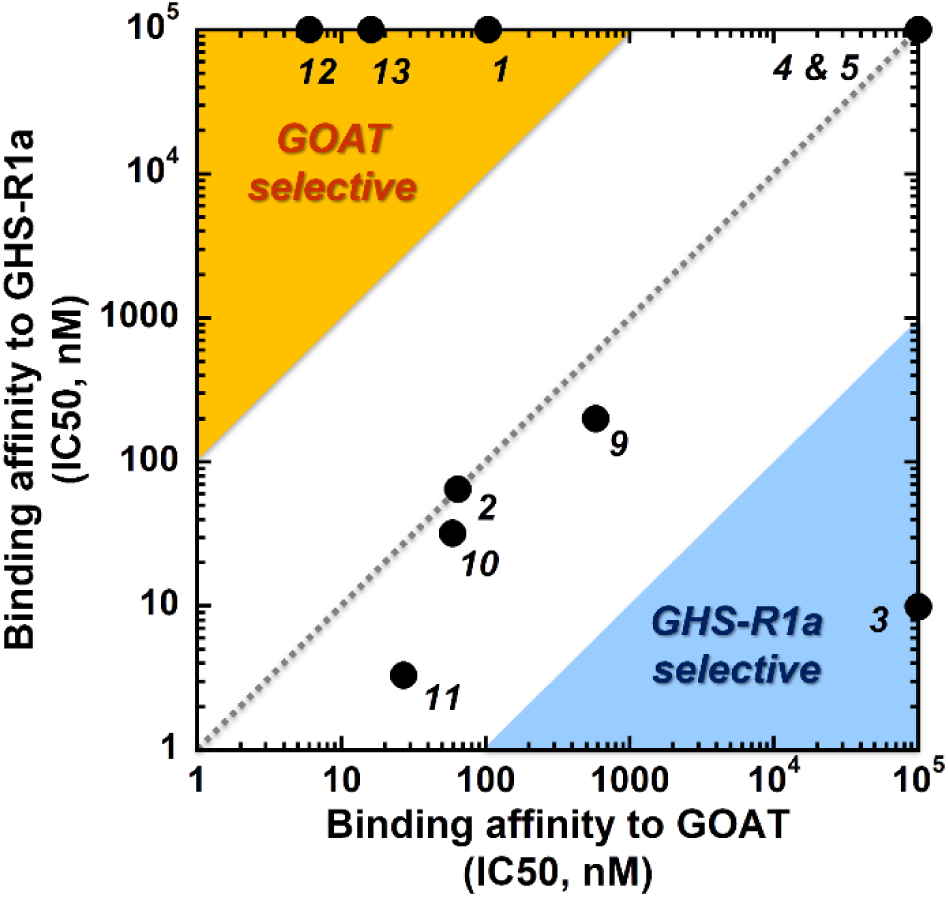
Ligand binding selectivity for GOAT and GHS-R1a. Binding affinities of each ligand for GOAT and GHS-R1a are plotted, with ligands in the upper left region (orange) exhibiting >100-fold selectivity for GOAT and ligand 3 in the lower right region (blue) displaying >100-fold selectivity for GHS-R1a binding. Ligand numbering and binding affinities (expressed as IC_50_ values) are provided from Table 1.

**Figure 2.**
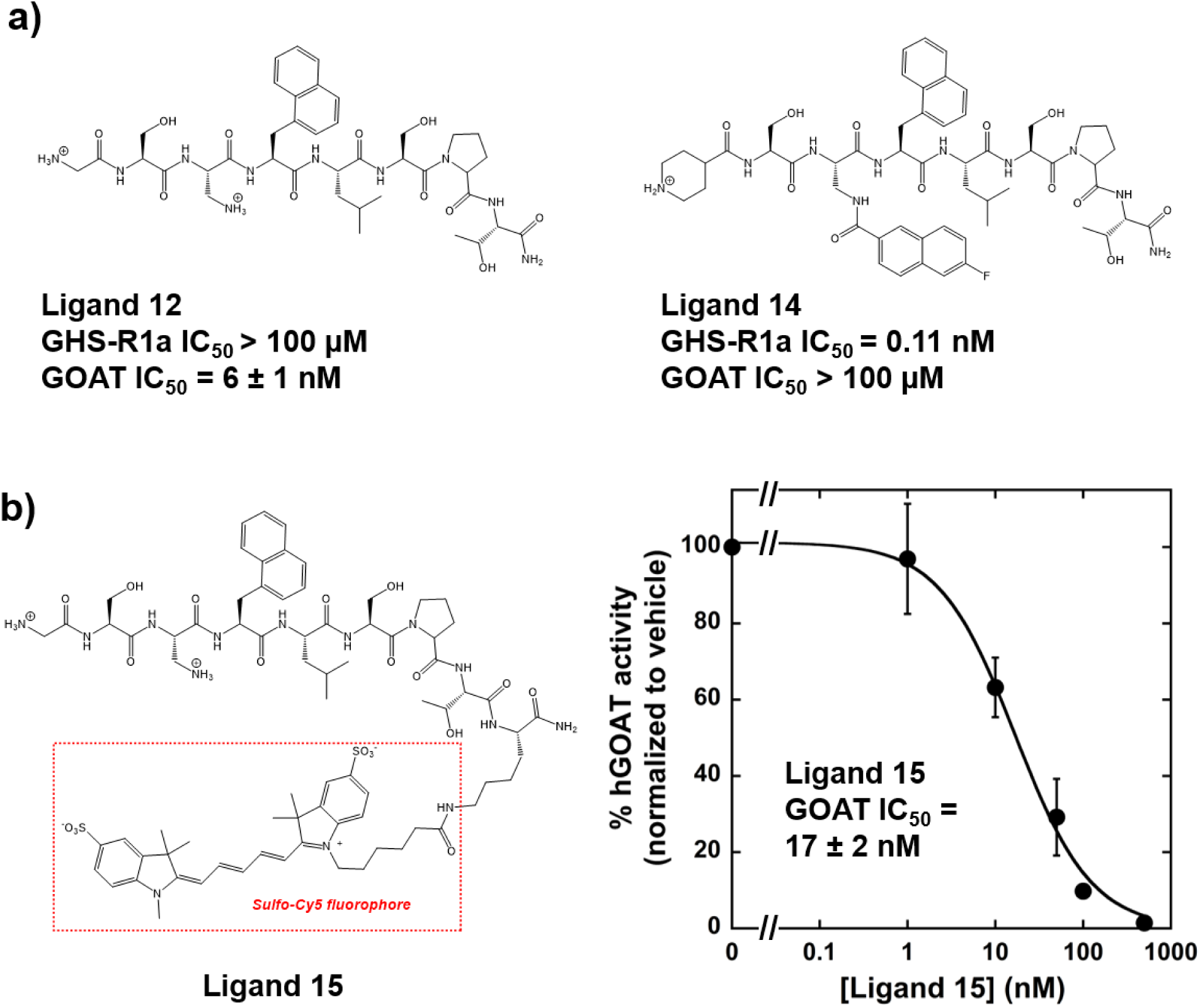
Orthogonal GHS-R1a and hGOAT ligands and fluorescently labeled specific GOAT ligand 15. a) Structures of the most potent and specific ligands for GHS-R1a and GOAT, with IC_50_ values measured against both ghrelin binding proteins; IC_50_ value for ligand **14** against GHS-R1a cited from Charron et al.^22^ b) Fluorescent ligand designed for strong binding to hGOAT. Structure of H-GS-Dap-Nal-1-LSPTK(SulfoCy5)-NH_2_ (ligand **15**) containing a Sulfo-Cy5 fluorophore appended to a lysine in position 9 of the ghrelin analogue sequence, and inhibition of hGOAT activity by ligand **15** demonstrating IC_50_ = 17 +/− 2 nM. All reported IC_50_ values against hGOAT represent the average of three independent trials, and error bars represent one standard deviation.

### Ligand binding and uptake by human cells transfected with hGOAT

To demonstrate interaction between ligand **15** and hGOAT in a biologically relevant cellular setting, we transiently transfected HEK 293 cells with either a FLAG-tagged hGOAT construct or empty vector and imaged cells to determine hGOAT expression and ligand **15** binding. Following live cell incubation with the fluorescent ligand **15,** cells were washed, fixed and labeled with antibodies against FLAG. Using spinning-disk confocal microscopy, FLAG-hGOAT expressing cells were positive for ligand **15** Cy5 fluorescence with ligand fluorescence distributed throughout the cell whereas empty vector control expressing cells were not, consistent with GOAT expression being required for ligand binding and cellular uptake (Figure 3 and Supporting Table S2). In contrast, cells transfected with empty vector excluded ligand **15** demonstrating that this peptide is not intrinsically cell permeable or nonspecifically associating with cellular membranes.

**Figure 3.**
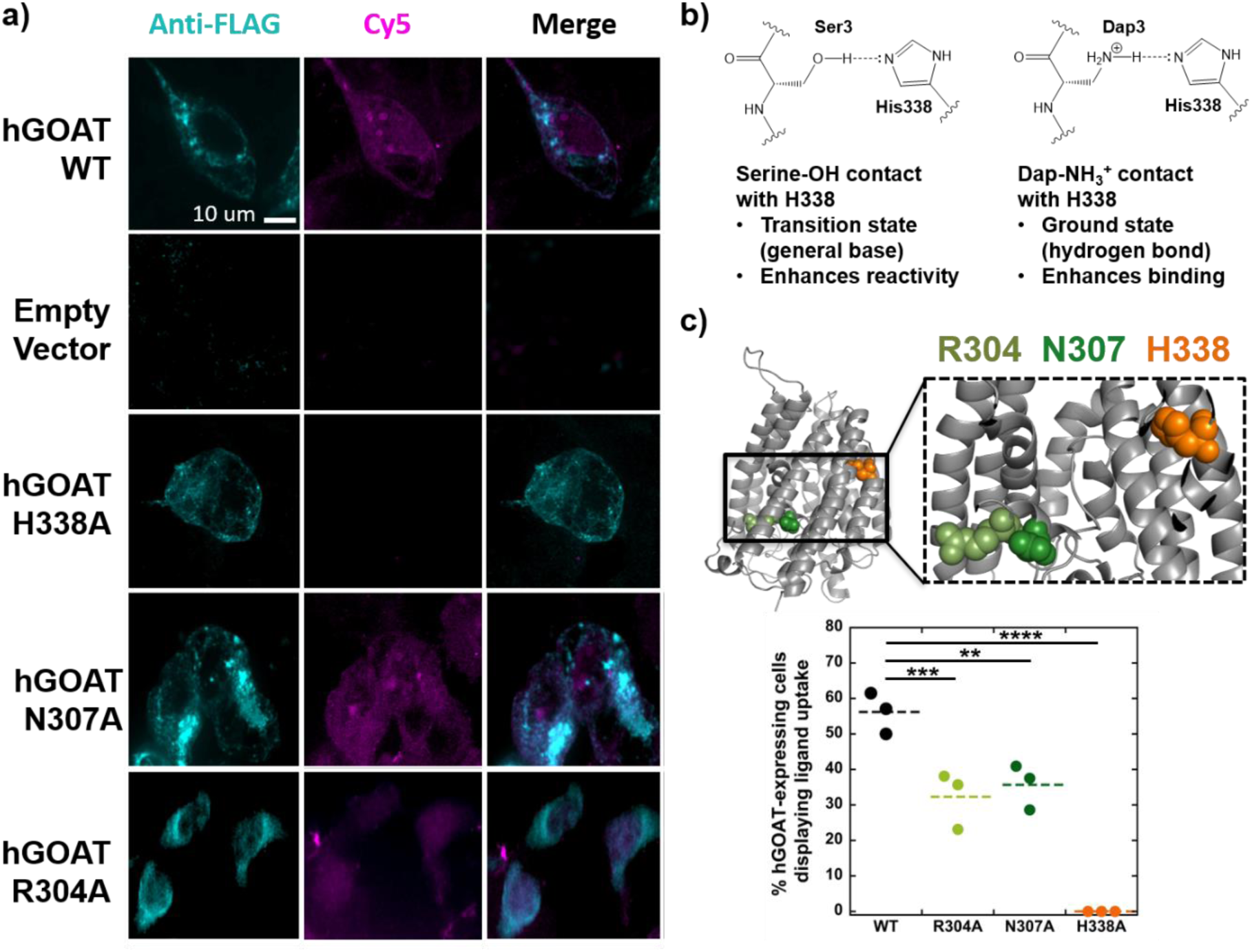
Interaction between the Dap amino group and H338 is essential for ligand uptake by GOAT-expressing cells. a) HEK 293 cells were transfected with empty vector or FLAG-tagged hGOAT/ mutant hGOAT as indicated. At 40 hrs post-transfection, cells were incubated with Cy5-ghrelin ligand **15** (purple) for 30 min and immunostained with anti-FLAG (cyan) for GOAT expression detection. Wild type, N307A, and R304A hGOAT expressing cells showed binding and internalization of ligand **15**. Empty vector and H338A hGOAT variants do not support peptide uptake. Scale bar, 10 µm. b) Proposed model for the interaction between the ghrelin serine 3 sidechain hydroxyl group and the ligand **15** Dap sidechain amine with H338 leading to general base catalysis and tighter ligand binding, respectively. c) Quantification of percent hGOAT-positive cells showing uptake of ligand **15** (n =100 cells, over n = 3 experiments) for wild type, R304A, N307A, and H338A hGOAT variants. The locations of R304 (olive green), N307 (forest green), and H338 (orange) are shown within the hGOAT structure. Dotted lines denote the average percent of each group. One-way ANOVA comparison test between wild type and hGOAT variants indicated significance with an adjusted P values of <0.01 (**, N307A), <0.001 (***, R304A), and < 0.0001 (****, H338A).

The uptake of ligand **15** by hGOAT-expressing cells provides the opportunity to define molecular interactions between the ligand and hGOAT responsible for ligand binding affinity. For example, the enhancement in binding to hGOAT upon amine substitution at the acylation site could arise from formation of a ground state hydrogen bond to an enzyme side chain which normally serves as a general base for serine acylation (Figure 3b). To explore ligand-enzyme interactions required for binding and cellular uptake, we have introduced three alanine mutations to functionally essential amino acids (Figure 3c). These three mutations all result in complete loss of hGOAT enzyme activity, but this loss of activity reflects interference in contributions by these residues at different steps of the hGOAT catalytic cycle.^29^ Histidine 338 is an absolutely conserved histidine in MBOAT family members and is proposed to serve as a general base interacting with the serine hydroxyl group.^26, 29^ We propose an interaction between the Dap side chain amine and H338 is partially responsible for the tight binding observed for Dap-containing peptide ligands to hGOAT. In contrast, both arginine 304 and asparagine 307 form interactions within the octanoyl-CoA acyl donor-binding site, which would not directly impact ghrelin-mimetic peptide binding to hGOAT (Figure 3 and Supporting Table S2).

The H338A hGOAT variant does not maintain cell uptake of ligand **15** when expressed in HEK293 cells, supporting that an interaction between the Dap amino group and H338 is required for tight ligand binding. Furthermore, the loss of ligand uptake in the presence of H338A hGOAT expression argues against general loss of cell membrane integrity as the mechanism for ligand internalization. Given the integral membrane nature of GOAT,^19, 29^ we considered it possible that hGOAT expression may destabilize or disrupt membrane integrity, which could allow ligand cell penetration without requiring direct binding to hGOAT. In contrast to the H338A variant, both R304A and N307A variants supported ligand **15** internalization as expected for mutations predicted to compromise the octanoyl-CoA binding pocket but not the ghrelin binding site within the hGOAT catalytic channel.

### Endogenous GOAT expression in prostate cancer cells supports ligand uptake

While unlikely, it is possible the hGOAT interaction with extracellular peptides in transfected HEK 293 cells reflects aberrant trafficking to the plasma membrane resulting from enzyme overexpression. To complement ligand uptake studies using hGOAT overexpression in transfected HEK293 cells, we examined human cell lines with endogenous GOAT expression for similar ligand binding and uptake. For these studies, we utilized LNCaP and 22Rv1 prostate cancer lines which have been reported to overexpress GOAT.^14–15, 30^ Immunofluorescence imaging using an anti-MBOAT4 antibody revealed robust hGOAT expression in both prostate cancer lines (Figure 4a-b and Supporting Figures S3 and S4). To determine whether a fraction of endogenous hGOAT is exposed on the plasma membrane to the extracellular environment, immunofluorescent labeling was repeated in 22Rv1 cells under non-permeabilizing conditions and compared to labeling with standard cell permeabilization. Non-permeabilizing conditions yielded hGOAT labeling at the cell surface with absence of immunofluorescence within the cell (Figure 4c) consistent with hGOAT being accessible to extracellular antibodies at the cell surface.

**Figure 4.**
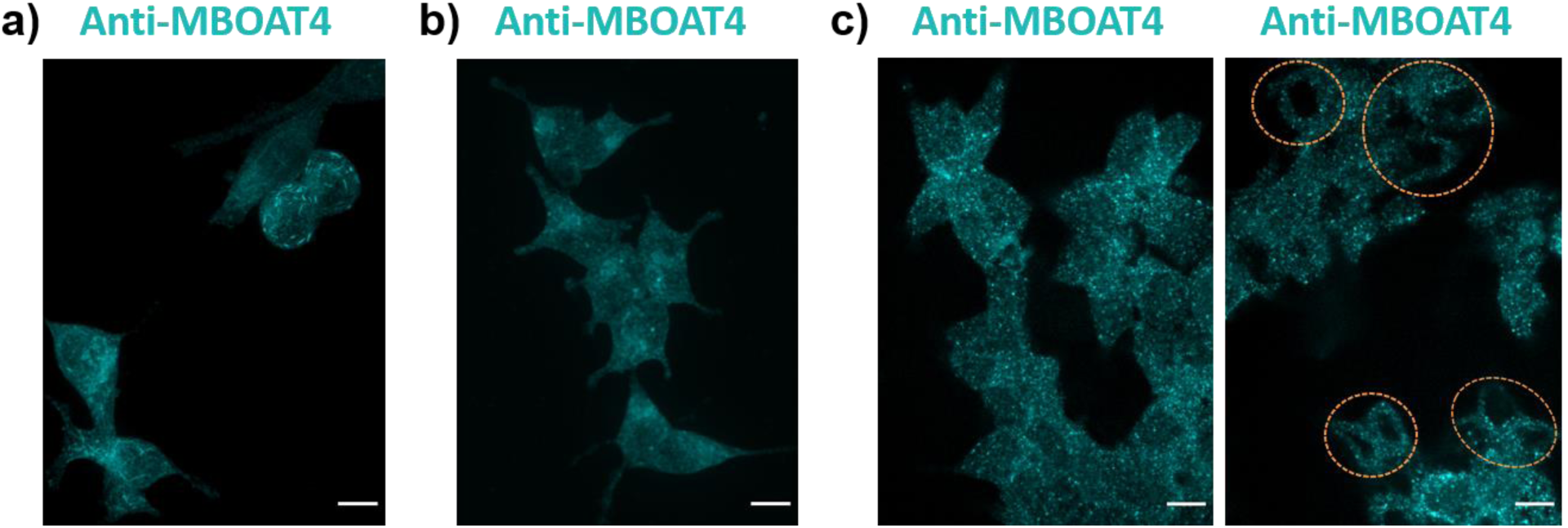
GOAT expression in prostate cancer cells leads to exposure to the extracellular space. a) GOAT imaged in LNCaP PCa cells by anti-MBOAT4 immunofluoresence. b) GOAT imaged in 22Rv1 PCa cells by anti-MBOAT4 immunofluoresence. c) Immunofluoresence imaging of GOAT in 22Rv1 cells under permeabilizing (left) and non-permeabilizing (right) conditions indicates GOAT can interact with antibodies outside the cell; cells whose interiors lie on the confocal plane are circled in orange. Cells were prepared for imaging as described in the Experimental Procedures. Scale bars, 10 μm.

We next examined binding and cellular uptake of ligand **15** in prostate cancer cells expressing hGOAT to compare with our earlier studies of hGOAT-transfected cells. Confocal imaging confirmed ligand binding and uptake in both 22Rv1 and LNCaP cells with 100% of the prostate cancer cells exhibiting ligand binding (Figure 5a-c). Ligand binding and uptake became more pronounced at higher ligand concentrations (Figure 5d). Labeling of 22Rv1 PCa cells with ligand **15** was significantly less efficient at 4 °C than at 37 °C consistent with ligand binding and uptake requiring unimpeded plasma membrane trafficking for ligand internalization (Figure 5e). Selective uptake of ligand **15** by hGOAT was probed by treating cells with an unlabeled competitor ligand to saturate cell surface-exposed hGOAT to block ligand **15** binding. Co-incubation with a large excess of unlabeled competitor ligand **12** similarly led to a significant reduction of ligand **15** internalization consistent with specific ligand recognition and uptake mediated through ligand binding to cell surface exposed hGOAT (Figure 5f-g).

**Figure 5.**
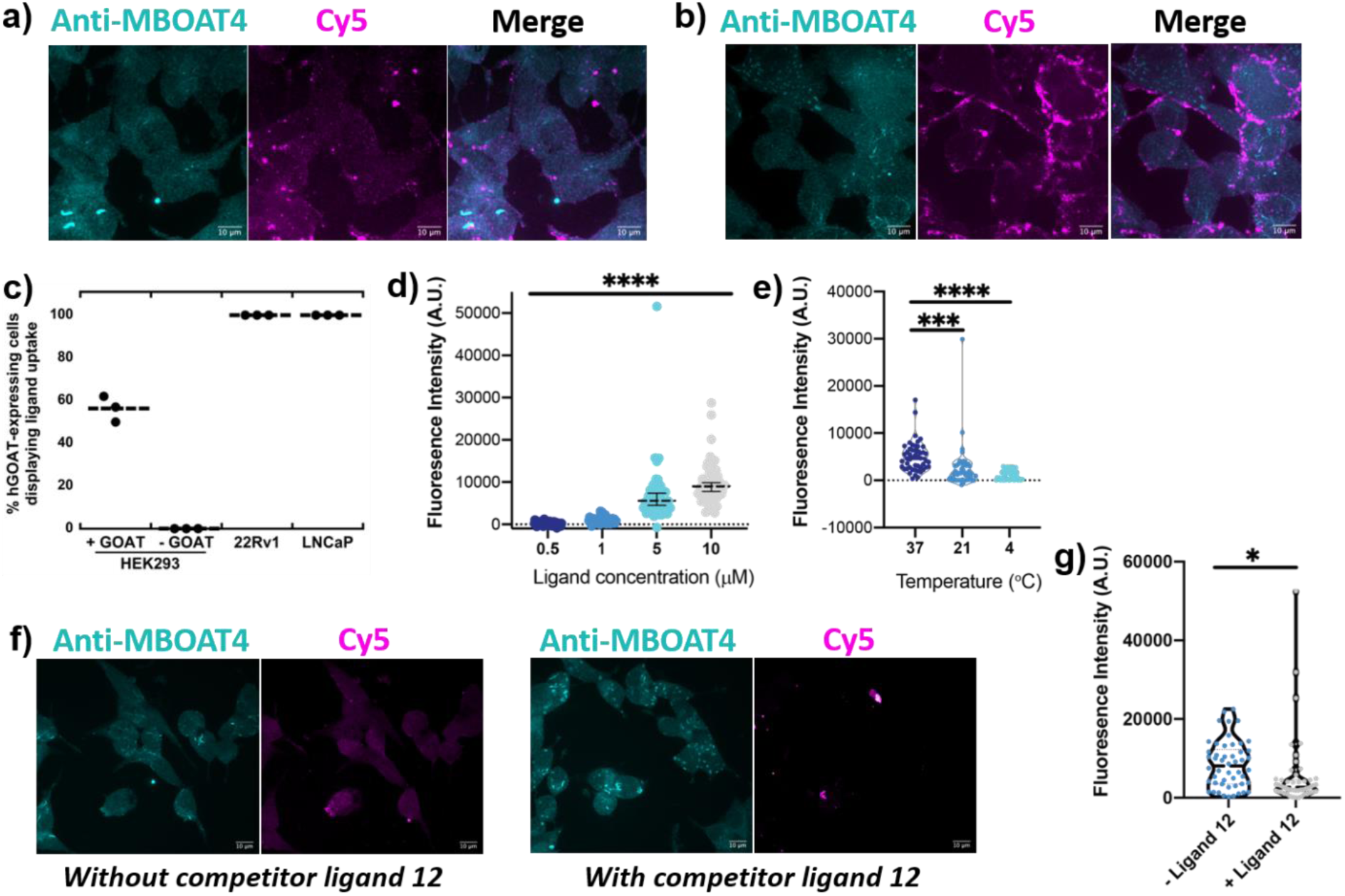
Endogenous hGOAT expression in prostate cancer cells supports GOAT ligand binding and uptake. a) 22Rv1 cells were incubated with Cy5-ghrelin ligand **15** (purple) for 30 min and immunostained with anti-MBOAT4 (cyan) for GOAT expression detection. b) LNCaP cells were incubated with Cy5-ghrelin ligand **15** (purple) for 30 min and immunostained with anti-MBOAT4 (cyan) for GOAT expression detection. c) Quantification of cells showing uptake of ligand **15** (n =50 cells, over n = 3 independent experiments) for HEK 293, HEK 293 transfected with hGOAT, 22Rv1, and LNCaP cells. d) Dependence of ligand **15** uptake by 22Rv1 cells on ligand concentration in the cell media, as reflected by total cell fluorescence. One-way ANOVA comparison tests between all ligand concentrations indicated significance with adjusted P values of <0.0001 (****). e) Temperature dependence of ligand **15** uptake in 22Rv1 cells. One-way ANOVA comparison tests between 37 degrees, 21 degrees, and 4 degrees indicated significance at both lower temperatures with adjusted P values of < 0.001 (***, 37 vs 21) and <0.0001 (****, 37 vs 4). f) Incubation of 22Rv1 cells and ligand **15** in the presence of unlabeled competitor ligand **12** reduces ligand **15** uptake. g) Quantification of ligand **15** uptake in 22Rv1 cells in the absence and presence of a competitor ligand, as indicated by total cell fluorescence. The reduction in presence of the competitor ligand was determined to be significant with p < 0.05 (*) by an unpaired T-test. Dotted lines denote the average values for each data group. Scale bars, 10 μm.

Originally assigned as an ER-resident enzyme responsible for acylating ghrelin during hormone maturation prior to secretion,^7^ our work provides the first direct evidence for GOAT exposure to the extracellular space and interaction with soluble peptides. These studies were enabled by the creation of a specific ghrelin-mimetic ligand for GOAT, which allows for direct detection and investigation of this ghrelin-binding protein without interference from the GHS-R1a receptor. Our development of ghrelin-based ligands opens the door for creating non-invasive imaging agents targeting GOAT, while also providing the first functional connection between the active site of GOAT and ghrelin through the Dap amine-H338 interaction. Most unexpectedly, our studies indicate GOAT can bind extracellular peptides and facilitate cellular uptake.

The absolutely conserved histidine residue that serves as one of the defining characteristics of MBOAT family members (H338 in GOAT) has been suggested to act as a general base in the acylation reactions catalyzed by these enzymes.^19, 23, 26, 29, 31–42^ While this catalytic role has not been conclusively demonstrated in any MBOAT, the dependence of ligand **15** uptake on the presence on H338 in GOAT supports a direct interaction between the acylation site serine in ghrelin and this conserved histidine (Figure 3b). The enhanced binding affinity of ghrelin ligands with amine modifications at the serine acylation site likely arises from re-apportionment of the transition-state stabilization energy from the serine – histidine hydrogen bond/general base interaction during acyl transfer to ground-state binding enhancement from the amine/ammonium – histidine interaction in ligand **15**. This interaction provides the first functional connection between a residue within ghrelin and the GOAT active site, which is essential for further modeling of this enzyme and its catalytic architecture.^29^ Looking beyond GOAT, it will be interesting to examine similar hydroxyl to amine acylation site substitutions in other MBOAT substrates to determine if this simple atomic substitution provides a facile family-wide strategy for generating potent MBOAT inhibitors and identifying catalytic interactions in these enzymes.

Cellular uptake of ligand **15** requires a subpopulation of GOAT to be exposed on the plasma membrane where it can bind ghrelin and desacyl ghrelin from outside the cell, consistent with detection of GOAT in intracellular and plasma membranes of lipid trafficking vesicles in blood marrow adipocytes using immunogold staining.^20^ Our unambiguous demonstration of ligand binding and cellular uptake supports an expanded view of GOAT’s involvement within the ghrelin trafficking and signaling pathway in cells expressing GOAT (Figure 6). Our finding provides key support for a new branch of the ghrelin signaling pathway involving local re-acylation of desacyl ghrelin by plasma membrane exposed GOAT, which could provide a mechanism explaining the biological impact of treatment with desacyl ghrelin.^7, 20–21, 43–46^ Ghrelin reacylation would allow cells and tissues presenting both GOAT and GHS-R1a on their surfaces to detect the total ghrelin level in circulation rather than only the acylated portion of the ghrelin pool in the bloodstream. Cellular integration of the “total ghrelin” signal could provide a readout for chronic organismal stress, whether from metabolic factors or other environmental stressors, rather than the acute signal provided by the rise and fall of the hydrolytically susceptible pool of ghrelin secreted from the gastrointestinal tract.^47–50^ This two-factor signaling system may provide mechanisms to explain the complex and sometimes paradoxical biological signaling behavior observed for ghrelin and desacyl ghrelin, essential features of this endocrine system that remain to be fully understood.

**Figure 6.**
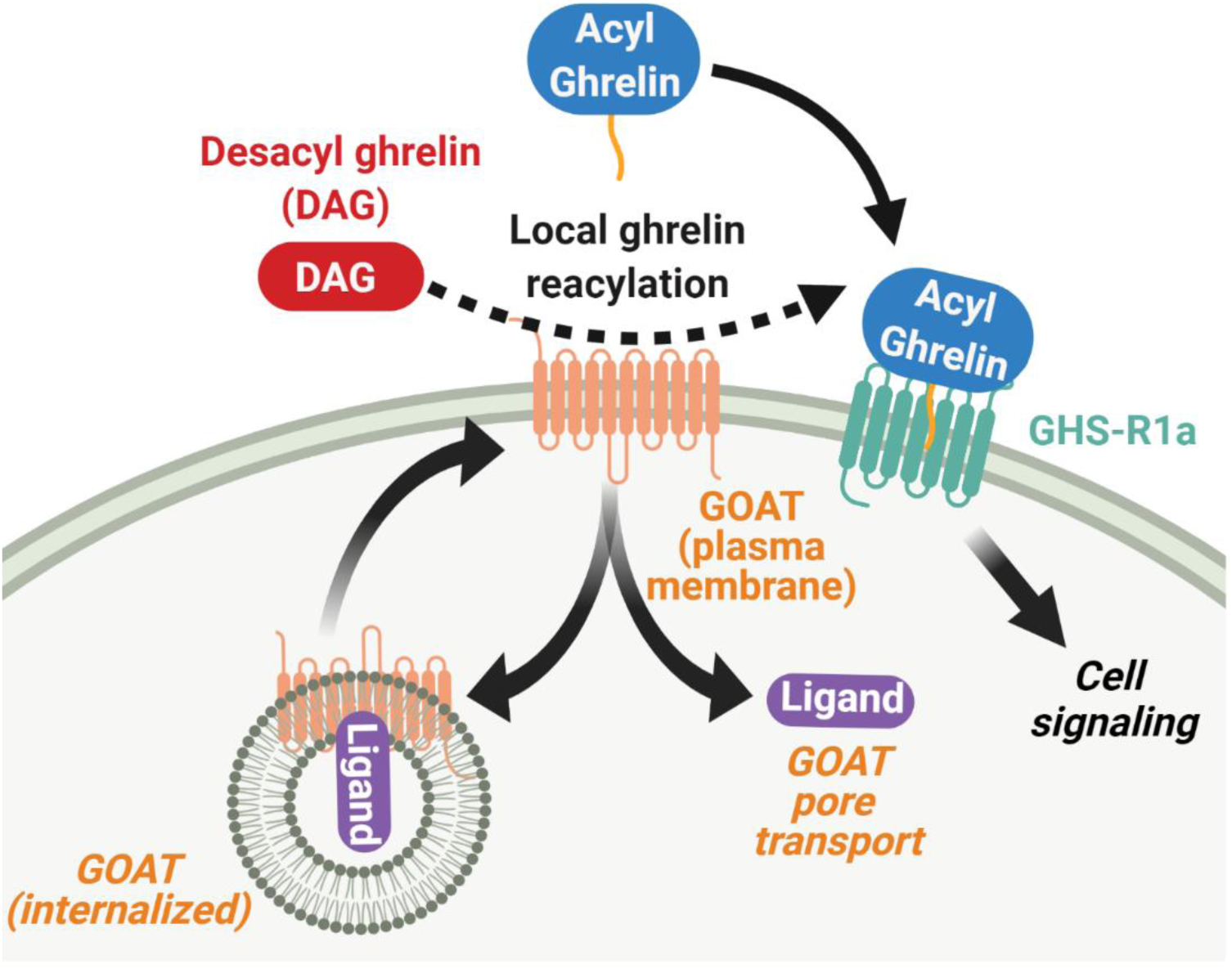
An expanded view of GOAT cellular localization and signaling roles. In cells presenting GOAT on their plasma membrane, GOAT could catalyze reacylation of desacyl ghrelin which can then activate GHS-R1a signaling. GOAT can also facilitate cellular uptake of peptide ligands through either enzyme-ligand complex internalization or ligand transport through the GOAT internal channel. Figure created using BioRender.

Given GOAT overexpression reported in prostate and breast cancer,^14–18, 30^ the role of surface-exposed GOAT in ghrelin signaling in cancer cells and the utility of GOAT as a cancer biomarker represent compelling new areas for investigation. Validating GOAT as a cancer biomarker will require GOAT expression analysis and ligand binding/uptake studies in related non-cancerous cells, such as the RWPE-1 cell line for applying GOAT as a prostate cancer biomarker.^14, 30^ In cancer cell lines and tissues found to overexpress GOAT, defining the impact of this overexpression on cell signaling is the next step in understanding how these transformed cells are affected by organismal metabolic and endocrine states as reflected by ghrelin and des-acyl ghrelin concentrations. We anticipate applying selective GOAT ligands, GOAT localization analysis, and ghrelin-dependent cell signaling studies in future work to explore this compelling intersection of ghrelin signaling and cancer biology.^13, 51^

Ligand internalization can be facilitated by GOAT through at least two distinct mechanisms: (1) GOAT-ligand complex transportation into the cell through membrane trafficking or (2) ligand transit into the cell by traversing GOAT as if the enzyme were a pore. In mechanism (1), the GOAT-ligand complex would be internalized by endocytosis similarly to receptor-mediated ligand uptake. In contrast, mechanism (2) allows GOAT to act as a ligand transporter at the plasma membrane without enzyme internalization. Computational modeling of GOAT supports the presence of an internal transmembrane channel that could act as a pore,^29^ and similar channels are observed in the crystal structure of the bacterial MBOAT DltB, the cryo-EM structures of Hhat, and structural models of PORCN.^32, 52–55^ We note our imaging studies do not support colocalization of ligand **15** with GOAT in either transfected HEK 293 cells or prostate cancer cells, which could potentially support ligand transport (mechanism 2) rather than enzyme-ligand internalization (mechanism 1). However, our studies were performed with fixed cells and the short length and small number of amine groups within the peptide ligand may lead to inefficient ligand crosslinking during fixation allowing intracellular dispersion during sample preparation. Further studies of GOAT and GOAT-ligand cellular trafficking will determine which of the proposed mechanisms is responsible for ligand internalization.

Looking to the future, we will investigate the mechanism and biological impact of peptide internalization by GOAT using the ligands reported in this work and expand our understanding of GOAT trafficking and localization within mammalian cells. Whether through transport of the GOAT:ligand complex by membrane trafficking or GOAT serving as a pore/transporter using its internal channel, this process presents a new and unanticipated function for integral membrane acyltransferases and may provide a novel avenue for intracellular drug/cargo delivery targeting cells expressing GOAT. We also suggest this work argues for examining the potential involvement of other integral membrane enzymes currently proposed to perform exclusively intracellular roles in extracellular interactions, chemistry, and signaling.

## Methods

### Peptide synthesis and characterization

Details of peptide synthesis and characterization are provided in the Supporting Information.

### GHS-R1a receptor binding assays

Peptide binding affinity for the ghrelin receptor was determined using a competitive radioligand-displacement binding assay,^22^ with details provided in the Supporting Information.

### hGOAT inhibition assays

Assays were performed using previously reported protocols,^56–57^ as described in the Supporting Information.

### Construction of hGOAT WT and mutants

Site-directed mutagenesis was performed on pcDNA 3.1 (+) mammalian expression vector containing a hGOAT insert cloned from our previously reported pFastBacDual vector (Invitrogen) using the *EcoR*I and *Xba*I restriction sites, resulting in the pcDNA3.1_Mb4.WT (hGOAT) construct.^56^ This construct contains a C-terminal FLAG epitope tag, a polyhistidine (His6) tag, and 3x human influenza hemagglutinin (HA) tags appended downstream of a TEV protease site.^57^

### hGOAT transfection in HEK 293 cells

Mammalian cell line HEK 293 (ATCC) was maintained in 75 mL vented tissue culture flasks (Celltreat) and kept to 70% confluency before splitting. All cells were cultivated in complete DMEM (Dulbecco’s Modified Eagle’s Medium supplemented with 10% fetal bovine serum (FBS) and 1% (v/v) penicillin-streptomycin (MediaTech) in a humidified atmosphere with 5% CO_2_ at 37°C. For transfection of WT hGOAT, mutant hGOAT, and empty vector (EV) cells were plated at a density of 1 × 10^6^ per well in 2 mL of complete DMEM in a 6-well plate per well (Corning) with sterile 12 mm 1.5 glass coverslips in each well (Warne Instruments). The cells were incubated for 16 hours prior to transfection. The DNA-transfection reagent complex was prepared by combining 4 µg of pcDNA3.1_Mb4.WT (hGOAT) or mutant plasmid and 9 µL Lipofectamine 2000 transfection reagent (Invitrogen) in a total volume of 500 µL supplement free DMEM followed by incubation for 30 min at room temperature. The cells were then transfected with the DNA-transfection reagent complex by drop wise addition into the plate wells.

### GOAT ligand labeling and immunofluorescence imaging in HEK293 cells

Following transfection for 40 hours, coverslips with attached cells were removed from the wells, washed with 1x phosphate-buffered saline (PBS) (Cellgro), and incubated with 10 µM ligand **15** (500 µL/well) for 30 minutes at 37°C. Following washing with 1x PBS, cells were fixed with 4% paraformaldehyde for 20 minutes at room temperature, washed with 1x PBS, quenched in 50 mM NH_4_Cl for 10 minutes at room temperature, and washed with 1x PBS. For antibody staining, all steps were performed in a Parafilm dark chamber at room temperature (unless otherwise specified) with humid atmosphere. Cells were blocked with PBSAT buffer (PBS + 1% Bovine serum albumin, 0.1% triton) for 30 minutes in the dark chamber, followed by aspiration of buffer without allowing the coverslip to dry. Cells were then incubated with primary antibody Rabbit anti FLAG (DYKDDDDK) antibody (Sigma, F7425) diluted in 1x PBS overnight in the dark chamber at 4°C. The following day, cells were washed with 1x PBS and incubated with secondary antibody Alexa Fluor 488-conjugated goat anti-rabbit (Jackson Immuno Research, 709-545-149) for 1 hour. Following antibody incubations, cells were washed three times, mounted on slides with DAPI, and analyzed by confocal microscopy. Images were taken on a Leica DMi8 STP800 (Leica, Bannockburn, IL) equipped with an 89 North–LDI laser with a Photometrics Prime-95B camera taken with a Crest Optics: X-light V2 Confocal Unit spinning disk. Optics used were HC PL APO 63×/1.40 NA oil CS2 Apo oil emersion objective.

### Prostate cancer cell line culture

Prostate cancer (PCa) cells LNCaP (ATCC, CRL-1740) and 22Rv1 (ATCC, CRL-2505) were maintained in 75 mL vented tissue culture flask (Celltreat) at 37°C, 5% CO2 in complete 1X RPMI (Roswell Park Memorial Institute) media supplemented with 10% FBS and 1% (v/v) penicillin-streptomycin (Mediatech). Cell lines were passaged upon reaching 70% confluency (∼2-3 days). For microscopy studies, PCa cells were plated in 2 mL of complete 1X RPMI in a 6-well plate (Corning) containing a sterile 12 mm poly-L-lysine coated glass coverslips (Neuvitro, GG-12-PDL) in each well. The cells were allowed to adhere and reach 70% confluency on coverslips prior to labeling.

### Immunofluorescence staining of hGOAT in PCa cell lines

Confluent PCa cells grown on coverslips were fixed with 4% paraformaldehyde for 20 minutes at room temperature and quenched in 50 mM NH4Cl for 10 minutes at room temperature. For antibody staining, all steps were performed in a Parafilm dark chamber at room temperature with humid atmosphere. Cells were blocked under either permeabilizing conditions with PBSAT buffer or under nonpermeabilizing conditions with PBSA (PBS + 1% Bovine serum albumin) for 30 minutes in the dark chamber. Cells were then incubated with 1:80 Rabbit anti MBOAT4 polyclonal antibody (Cayman, #18614) for 1 hour in the dark chamber at 4°C and then cells were incubated with 1:1000 secondary antibody Alexa Fluor 488-conjugated goat anti-rabbit (Thermo, A21206) for 1 hour at room temperature. Following antibody incubations, cells were extensively washed, mounted on slides with Prolong containing DAPI (Thermo, P36971), and analyzed by confocal microscopy.

### GOAT ligand labeling and imaging in PCa cells

Upon reaching confluency, coverslips with attached PCa cells were treated with ligand at the indicated concentration for 30 min at room temperature (unless otherwise stated). Cells were washed 3x with 1x phosphate-buffered saline (PBS) (Cellgro) and then fixed for immunofluorescence staining as described above. For ligand competition experiments, cells were incubated with 5 µM ligand **15** alone, 5 µM ligand **15** with 40 µM ligand **12**, or PBS alone for 30 minutes at room temperature. For variable temperature experiments, PCa cells were preincubated in a tissue culture refrigerator (4 °C), on the benchtop (21 °C) or an incubator (37°C) for 30 min prior to addition of ligand **15**. The cells were further incubated with ligand **15** at those temperatures for 30 minutes and then processed for imaging as described above.

### Image analysis

The entire cell was imaged at 0.2-μm step-intervals and displayed as maximum projections (ImageJ). The fluorescence range of intensity was adjusted identically for each image series. Graphs and statistical analyses were completed using Graphpad Prism software, with specific tests and p-values provided in figure captions. All images were set to a resolution of 300 DPI or greater after image analysis from raw data. Total cell fluorescence was determined by calculating the integrated density of mean gray value in a cell area compared to background as shown in equations 4 and 5.

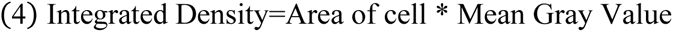

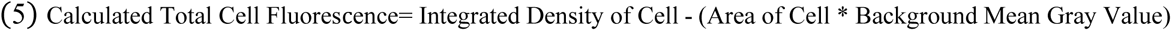

## Data availability

All data are contained in the manuscript and Supporting Information.

## Supporting information

This article contains supporting information: Supporting methods; Analytical data for peptide ligands; Dose-response curves for hGOAT inhibition by peptide ligands; Quantification of cell imaging analysis for hGOAT expression and peptide ligand uptake in transfected HEK 293 cells; Western blot confirmation of hGOAT expression in transfected HEK293 cells; Western blot validation of anti-MBOAT4 antibody; Validation of anti-MBOAT4 antibody IF labeling in HEK293 and prostate cancer cells.

## Acknowledgements

We gratefully acknowledge helpful discussions and suggestions from members of the Hougland, Hehnly, and Luyt research groups.

## Author Contribution

MBC, TRD, ER Cleverdon, HH, LGL, and JLH conceived of the project. MBC, TRD, ER Cleverdon, MB, NK, ER Curtis, MDC, MRP, SXN, YM-R, and MAS developed protocols and carried out experiments. MBC, TRD, MDC, HH, LGL, and JLH interpreted data and wrote the manuscript. JLH supervised the project.

## Funding and additional information

This work was financially supported by Syracuse University, the American Diabetes Association (grant #1-16-JDF-042 to JLH), and National Institutes of Health grants GM134102 (JLH), GM127621 (HH), and GM130874 (HH). This work was supported by the U.S. Army Medical Research Acquisition Activity through Prostate Cancer Research Programs under Award no. W81XWH2010585 (H.H.). TRD acknowledges support from the US Department of Education (GAANN Award P200A160193), and this manuscript is also based in part upon work supported by the National Science Foundation under Grant No. CHE-1659775. Funding was also provided by the Natural Sciences and Engineering Research Council (NSERC) of Canada to LL. MDC acknowledges support through the NSERC Postgraduate Scholarships-Doctoral Program and the Molecular Imaging Graduate Program of the University of Western Ontario. The content is solely the responsibility of the authors and does not necessarily represent the official views of the National Institutes of Health.

## Conflict of Interest

JLH has patent interests in ligands reported in this work.

## Abbreviations

GHS-R1a: growth hormone secretagogue receptor type 1a
GOAT: ghrelin *O*-acyltransferase hGOAT, human ghrelin *O*-acyltransferase
C8: eight-carbon fatty acid octanoate
ER: endoplasmic reticulum
Dap: diaminopropionic acid
PFPN: pentafluorophenyl naphthoate
Aib: aminoisobutyric acid
Inp: isonipecotic acid
Nal-1: 1-naphylalanine
Nal-2: 2-naphylalanine
AcDan: acrylodan
MAFP: methyl arachidonyl fluorophosphonate
DMSO: dimethyl sulfoxide
HEPES: 4-(2-hydroxyethyl)-1-piperazineethanesulfonic acid
Tris: 2-Amino-2-hydroxymethyl-propane-1,3-diol
HPLC: high performance liquid chromatography
MALDI: matrix-assisted laser desorption/ionization
IC_50_: half maximal inhibitory concentration
DMF: *N*,*N*-dimethylformamide
HCTU: *O*-(6-chlorobenzotriazol-1-yl)-*N*,*N*,*N*′,*N*′-tetramethyluronium hexafluorophosphate
DIPEA: *N*,*N*-diisopropylethylamine
TFA: trifluoroacetic acid
TIS: triisopropylsilane
TBME: *tert*-butyl methyl ether
DMEM: Dulbecco’s modified eagle’s medium
FBS: fetal bovine serum
WT: wildtype
EV: empty vector
PBS: phosphate-buffered saline
PBSAT: PBS + 1% bovine serum albumin-0.1% triton
SDS: sodium dodecyl sulfate
PVDF: polyvinylidene difluoride
TBST: tris 10 buffered saline, DAPI, 4,6-diamidino-2-phenylindole
HEK: human embryonic kidney fibroblasts.

## Notes

### Summary of Updates

This revision includes studies with cell lines endogenously expressing GOAT. These additional studies demonstrate GOAT ligand uptake and internalization occurs in both transfected cells and cells naturally expressing GOAT.

