## Supplemental Info for "A remarkably specific ligand reveals ghrelin *O*-acyltransferase interacts with extracellular peptides and exhibits unexpected cellular localization for a secretory pathway enzyme"

**Supporting Methods**

**Supporting Table S1**. Analytical data for peptide ligands

**Supporting Table S2.** Cell imaging and scoring for GOAT transfection and ligand **15** uptake.

**Supporting Figure S1.** Dose-response curves for hGOAT inhibition by peptide ligands

**Supporting Figure S2.** Western blot verification of hGOAT expression in transfected HEK 293 cells.

**Supporting Figure S3.** Western blot verification of anti-MBOAT4 antibody

**Supporting Figure S4.** Immunofluorescence validation of anti-MBOAT4 antibody in HEK293, 22Rv1, and LNCaP cells.

**Supporting Methods**

*Peptide synthesis and characterization.* Peptide synthesis was carried out using standard Fmoc solid-phase peptide synthesis on a Biotage SyroWave automated peptide synthesizer. Peptides were synthesized on a 0.1 mmol scale using Rink amide MBHA resin (0.39 mmol/g). The resin was initially swelled with dichloromethane (DCM), followed by Fmoc deprotection using 2 mL of 40% piperidine in *N*,*N*-dimethylformamide (DMF) for two cycles (3 min, 12 min). Amino acids were coupled with 4 equiv of Fmoc protected amino acid, 4 equiv. of *O*-(6-chlorobenzotriazol-1-yl)-*N*,*N*,*N'*,*N'*-tetramethyluronium hexafluorophosphate (HCTU), and 8 equiv of *N*,*N*-diisopropylethylamine (DIPEA) in *N*-methylpyrrolidinone (NMP). The mixture was added to the resin and vortexed for 40 min. These cycles were repeated until all amino acids were coupled to the resin.

Allyloxycarbonyl deprotection of diaminopropionic acid was performed manually under inert atmospheric $N_2$ conditions. DCM was dried over sieves for 24 h before adding 4 mL to the resin. An amount of 24 equiv of phenylsilane was then added to the peptide resin and shaken for 5 min. An amount of 0.15 equiv of tetrakis(triphenylphosphine)-palladium(0) was then added and allowed to react for 10 min. If desired, the resulting free amine was then acylated using 3 equiv of the corresponding acid (octanoic acid or 6-fluoro-2-naphthoic acid), 3 equiv of HCTU, and 6 equiv of DIPEA in DMF. The reaction mixture was left to couple overnight.

Full deprotection of the synthesized peptide was performed by adding a 2 mL mixture of 95% trifluoroacetic acid (TFA), 2.5% triisopropylsilane (TIS), and 2.5% water to the resin and allowing it to mix for 5 h. The cleaved peptide was precipitated from solution using ice-cold *tert*-butyl methyl ether (TBME) and centrifuged (3000 rpm, 10 min, 0 ˚C) resulting in a crude peptide pellet. The supernatant was decanted, and the resulting peptide pellet was dissolved in 20%

acetonitrile in water, frozen at −78 °C, and lyophilized to a white crude powder. Purification was performed using preparative HPLC-MS, and purity of the resulting peptides was analyzed using analytical HPLC-MS. These results are summarized in Supporting Table S1, with all compounds determined to have ≥95% purity.

*GHS-R1a receptor binding assays.* Peptide binding affinity for the ghrelin receptor was determined using a competitive radioligand-displacement binding assay.[1] Assays were performed using GHS-R1a transfected HEK 293 cells as the receptor source and human His[$^{125}$I]-ghrelin(1−28) (PerkinElmer Inc. NEX388010UC) as the radioligand. Human ghrelin(1−28) was used as a reference to ensure the validity of the results. A suspension of membrane from HEK 293/GHS-R1a cells (100,000 cells per assay tube) were incubated with ghrelin(1−8) peptide analogues (at concentrations of $10^{-6}$ M, $10^{-7}$ M, $10^{-8}$ M, $10^{-9}$ M, $10^{-10}$ M, $10^{-11}$ M, and $10^{-12}$ M) and His[$^{125}$I]-ghrelin (15 pM per assay tube) in binding buffer (25 mM HEPES, 5 mM magnesium chloride, 1 mM calcium chloride, 2.5 mM EDTA, and 0.4% BSA, pH 7.4). The resulting suspension was incubated for 20 min under shaking (550 rpm) at 37 °C. Unbound [$^{125}$I]-ghrelin was removed and the amount of [$^{125}$I]-ghrelin bound to the membranes was measured by γ counter. $IC_{50}$ values were determined by nonlinear regression analysis to fit a four-parameter dose−response curve using Prism 6 (version 6.0c). All binding assays were performed in triplicate.

*hGOAT inhibition assays.* Assays were performed using previously reported protocols.[2-3] For each assay, membrane fraction from Sf9 cells expressing hGOAT was thawed on ice and homogenized by passage through an 18-gauge needle 10 times. Assays were performed with 50 µg of membrane protein, as determined by Bradford assay. Membrane fraction was pre-incubated with 1 µM methyl arachidonyl fluorophosphonate (MAFP) and unlabeled peptide inhibitor or vehicle as indicated in 50 mM HEPES pH 7.0 for 30 minutes at room temperature prior to reaction initiation.[4] All reactions were initiated by the addition of 1.5 μM $GSSFLC_{AcDan}$ and 300 μM octanoyl CoA. Reactions were incubated for 30 minutes at room temperature under foil. All assays were stopped with the addition of 50 μL of 20% acetic acid in isopropanol, and solutions were clarified by protein precipitation with 16.7 μL of 20% trichloroacetic acid, followed by centrifugation (1,000 x *g*, 2 minutes). The supernatant was then analyzed using reverse-phase HPLC with fluorescence detection as previously described.[2-3] Peak integrations for both substrate and product peaks were calculated using Chemstation for LC (Agilent Technologies). Data reported are the average of three independent determinations, with error reported as standard deviation.

For determination of $IC_{50}$ values, the percent activity at each inhibitor concentration was calculated from HPLC integration data using equations 1 and 2: To determine an $IC_{50}$ value for a given inhibitor, the plot of % activity versus [inhibitor] was fit to equation 3, with % $activity_0$ denoting hGOAT activity in the presence of the vehicle alone. All reported $IC_{50}$ values represent the average of a minimum of three independent trials.

$$(1)\ \%\ \text{activity} = \frac{\%\ \text{peptide acylation in presence of inhibitor}}{\%\ \text{peptide acylation in absence of inhibitor}}$$

$$(2)\ \%\ \text{peptide octanoylation} = \frac{\text{Fluorescence of acylated peptide}}{\text{Total peptide fluorescence (acylated and non-acylated)}}$$

$$(3)\ \%\ \text{activity} = \%\ \text{activity}_0 * \left(1- \frac{[\text{inhibitor}]}{[\text{inhibitor}] + IC_{50}}\right)$$

*Confirmation of hGOAT expression in transfected HEK 293 cells by Western blot.* hGOAT transfected cells were harvested by treatment with 0.25% trypsin-EDTA at 37°C for 5 minutes. Cells were then transferred to an Eppendorf tube and collected by centrifugation at low speed at room temperature. The media was aspirated and cells were resuspended in 1x sample buffer (0.33 M Tris HCl, pH 6.8, 0.1 M SDS, 14% glycerol, and 0.5 M DTT) and 50 mM HEPES pH 7.0 in a total volume of 45 µL. Samples were heated to 50.2 °C for 5 minutes and then incubated at room temperature for 15 min prior to gel loading.[3] Samples were loaded onto a 12 % Tris-glycine SDS-polyacrylamide gel and run at 110 V for 1.5 hrs. Each gel was loaded with a negative control (empty vector (EV) microsomal protein) and amino-terminal FLAG-BAP Fusion protein as a positive control (Millipore Sigma, P7582-100UG, 1:200 dilution, 50 µL total volume) using a previously published protocol.[1] Following SDS-polyacrylamide electrophoretic separation, proteins were transferred to a polyvinylidene difluoride (PVDF) membrane (BioRad, Trans-Blot turbo RTA transfer kit). The PVDF membrane was activated with methanol incubation for 30 seconds followed by equilibration in transfer buffer (20% v/v methanol, 48 mM Tris base, 39 mM glycine and 0.034% v/v SDS) before transfer. Proteins were transferred to the membrane for 30 minutes at 1.3 A / 25 V using a transfer kit per manufacturer's instructions. Following transfer electroblotting, the PVDF membrane was blocked for 3 hours with 5% v/v nonfat milk in TBST buffer (Tris 10 buffered saline (TBS, 0.05M Tris and 0.14M NaCl) with 0.1% v/v Tween 20). FLAG antibody (HRP-conjugated DYKDDDDK Tag Antibody, Invitrogen catalog number PA1-984B-HRP, 1:2000 dilution, 10 mL total volume) was prepared in 5% nonfat milk in TBST buffer and membrane was incubated with the antibody overnight at 4 °C. The membrane was washed with TBST (6 x 5 mL) and treated with West Pico Chemiluminescent substrate-imaging reagent

(Thermo Scientific), followed by imaging on a ChemiDoc XRS+ gel documentation system (BioRad).

**Supporting Table 1**. Analytical data for peptide ligands

| Ligand | Peptide Sequence | $[M+H]^+$ calcd | $[M+H]^+$ found | Purity (%) | Yield (%) |
|---|---|---|---|---|---|
| 1 | H-GS-Dap-FLSPY-$NH_2$ | 855.4365 | 855.3387 | 99 | 36 |
| 2 | H-GS-(C8-Dap)-FLSPY-$NH_2$ | 981.5409 | 981.4415 | 98 | 20 |
| 3 | H-GS-(6FN-Dap)-FLSPY-$NH_2$ | 1027.4689 | 1027.5400 | 99 | 25 |
| 4 | H-Aib-S-Dap-FLSPY-$NH_2$ | 883.4678 | 883.3658 | 99 | 26 |
| 5 | H-Inp-S-Dap-FLSPY-$NH_2$ | 909.4834 | 909.4159 | 99 | 28 |
| 9 | H-GS-(C8-Dap)-FLSPE-$NH_2$ | 947.5202 | 947.8200 | 96 | 22 |
| 10 | H-GS-(C8-Dap)-FLSPN-$NH_2$ | 932.5205 | 932.8500 | 99 | 25 |
| 11 | H-GS-(C8-Dap)-FLSPT-$NH_2$ | 919.5253 | 919.8689 | 99 | 24 |
| 12 | H-GS-Dap-Nal-1-LSPT-$NH_2$ | 843.4365 | 843.3524 | 99 | 10 |
| 13 | H-GS-Dap-Nal-2-LSPT-$NH_2$ | 843.4365 | 843.3524 | 98 | 34 |
| 14 | H-Inp-S-(6FN-Dap)-Nal-1-LSPT-$NH_2$ | 1069.5159 | 1069.5227 | 99 | 33 |
| 15 | H-GS-Dap-Nal-1-LSPT-(SulfoCy5-K)-$NH_2$ | $[M+H]^{2+}$ = 798.3673 | $[M+H]^{2+}$ = 798.5341 | 95 | 7 |

**Supporting Table 2.** Cell imaging and scoring for GOAT transfection and ligand **15** uptake.

**Trial 1**

| **hGOAT variant** | **Total cells counted (n)** | **# Cells displaying hGOAT expression (anti-FLAG immunofluorescence)** | **# Cells displaying ligand 15 uptake (Cy5 fluorescence)** | **% of hGOAT-positive cells exhibiting ligand 15 uptake** |
|---|---|---|---|---|
| **Wild type** | 100 | 20 | 10 | 50 |
| **Empty Vector** | 100 | 0 | 0 | N/A |
| **H338A** | 100 | 13 | 0 | N/A |
| **R304A** | 100 | 26 | 6 | 23 |
| **N307A** | 100 | 28 | 8 | 29 |

**Trial 2**

| **hGOAT variant** | **Total cells counted (n)** | **# Cells displaying hGOAT expression (anti-FLAG immunofluorescence)** | **# Cells displaying ligand 15 uptake (Cy5 fluorescence)** | **% of hGOAT-positive cells exhibiting ligand 15 uptake** |
|---|---|---|---|---|
| **Wild type** | 100 | 21 | 12 | 57 |
| **Empty Vector** | 100 | 0 | 0 | N/A |
| **H338A** | 100 | 16 | 0 | N/A |
| **R304A** | 100 | 28 | 10 | 36 |
| **N307A** | 100 | 24 | 9 | 38 |

**Trial 3**

| **hGOAT variant** | **Total cells counted (n)** | **# Cells displaying hGOAT expression (anti-FLAG immunofluorescence)** | **# Cells displaying ligand 15 uptake (Cy5 fluorescence)** | **% of hGOAT-positive cells exhibiting ligand 15 uptake** |
|---|---|---|---|---|
| **Wild type** | 100 | 26 | 16 | 62 |
| **Empty Vector** | 100 | 0 | 0 | N/A |
| **H338A** | 100 | 10 | 0 | N/A |
| **R304A** | 100 | 21 | 8 | 38 |
| **N307A** | 100 | 22 | 9 | 41 |

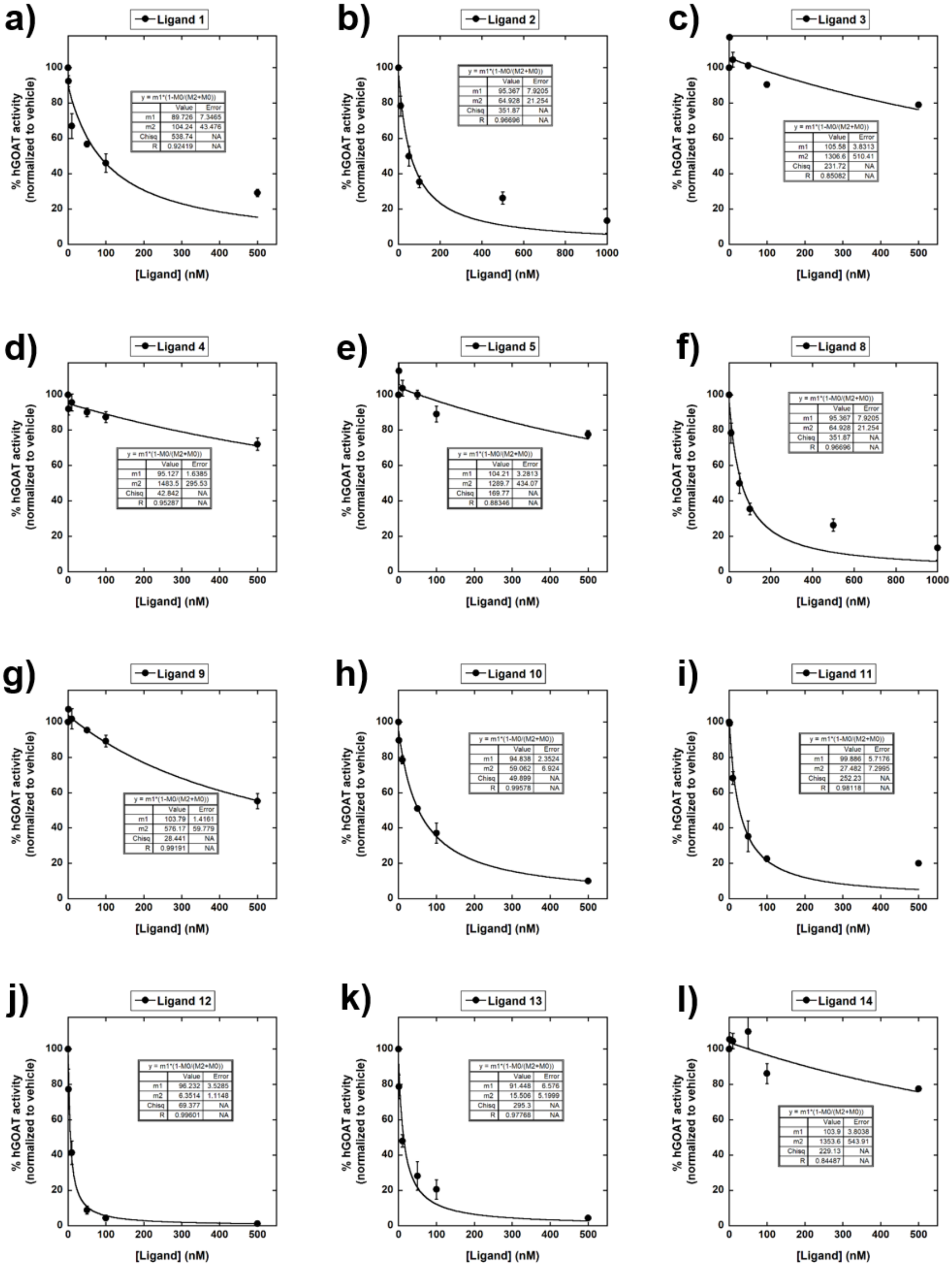


**Supporting Figure 1. Dose-response curves for hGOAT inhibition by peptide ligands.** All reported $IC_{50}$ values against hGOAT represent the average of three independent trials, and error bars represent one standard deviation. Inhibition was measured and analyzed as described in Experimental Procedures.

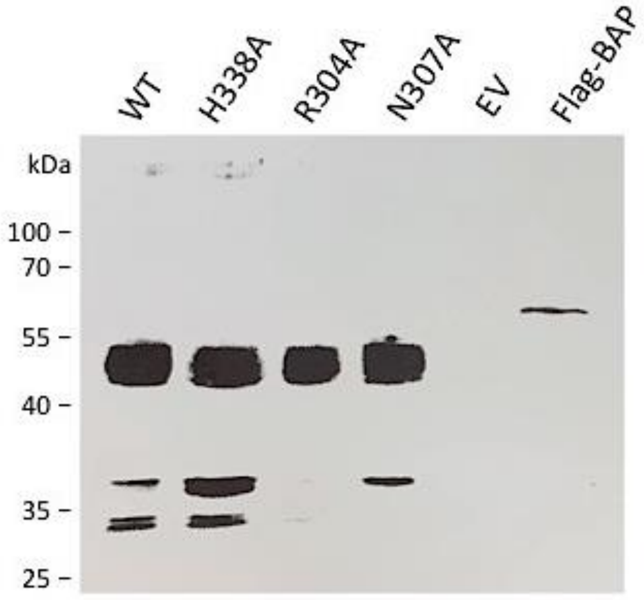


**Supporting Figure 2. Western blot verification of hGOAT expression in transfected HEK 293 cells.** hGOAT variant expression confirmation by anti-FLAG Western blotting with expected size of hGOAT variants (49 kDa). Western blots were performed as described in the Methods section. WT, wild type hGOAT; EV, empty vector expression; FLAG, FLAG-BAP fusion protein positive control.

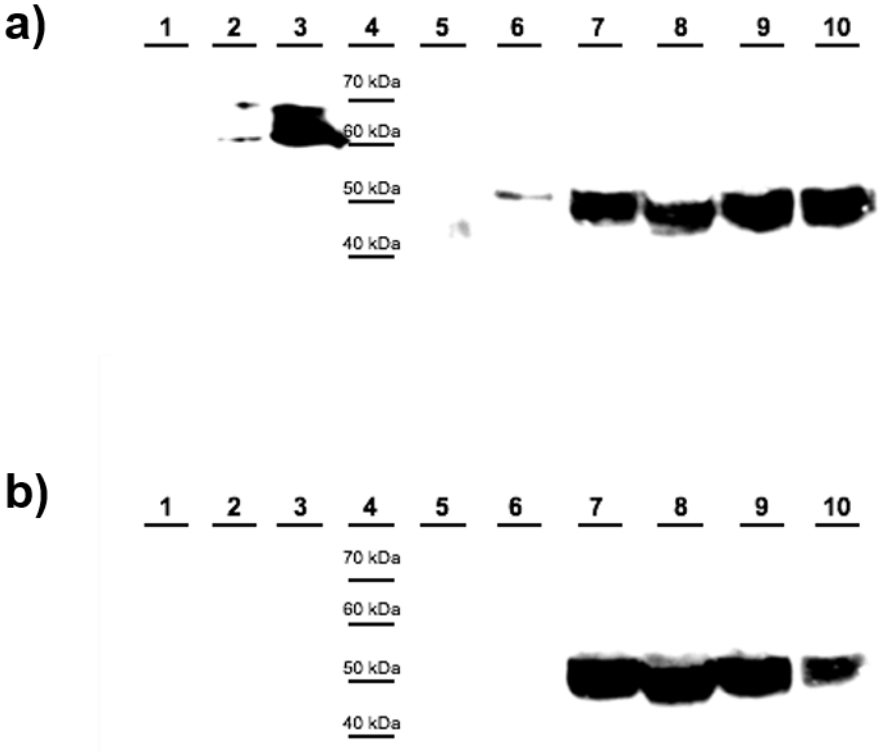


**Supporting Figure S3. Western blot verification of anti-MBOAT4 antibody.** Human and mouse ghrelin O-acyltransferases with a C-terminal 3xHA-Flag-$His_6$ tag were expressed in insect cells using the Bac-to-Bac baculoviral system (Invitrogen), and 30 μg total membrane protein was loaded in each well on a 10% SDS-PAGE gel. Protein was transferred to a PVDF membrane and blotted with anti-FLAG antibody (a) and anti-human MBOAT4 antibody (b). Lanes: 1, Uninfected Sf9 membrane protein fraction; 2, Empty vector infected SF9 membrane protein fraction; 3, FLAG-BAP fusion protein positive control; 4, Protein Ladder; 5, Blank lane; 6, Mouse ghrelin O-acyltransferase (mGOAT)-infected Sf9 membrane protein fraction; 7-10, Human ghrelin O-acyltransferase (hGOAT)- infected SF9 membrane protein fractions. The FLAG-tagged hGOAT was detected by both antibodies, while FLAG-tagged mGOAT is only detected by the anti-FLAG antibody consistent with the antibody specificity reported by the manufacturer (Cayman Chemical). Western blots were performed as described in the Experimental section.

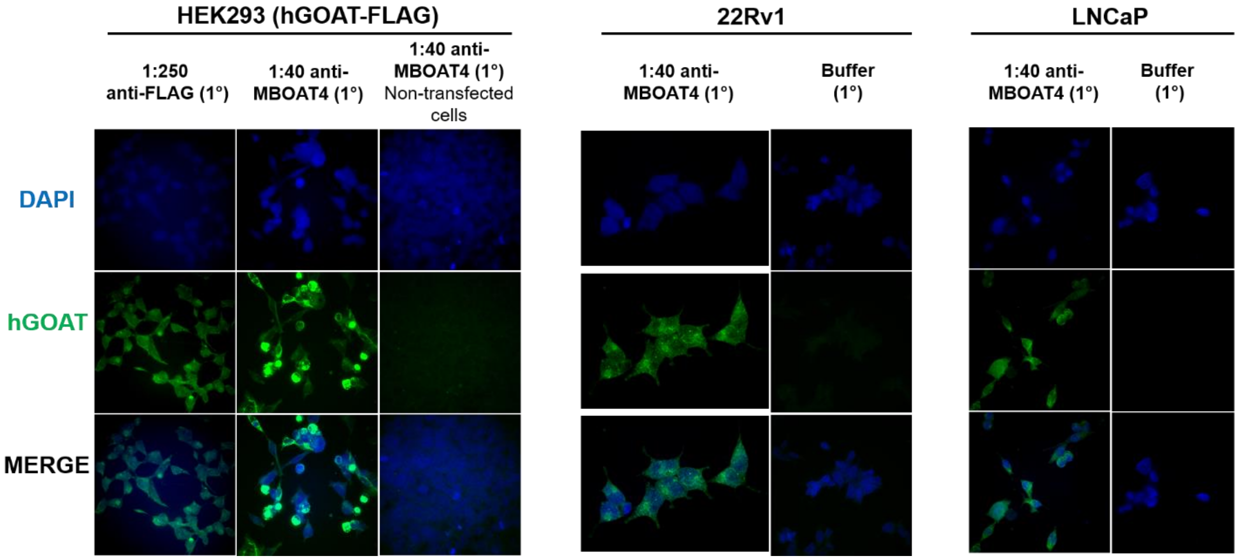


**Supporting Figure 4. Immunofluorescence validation of anti-MBOAT4 antibody in HEK293, 22Rv1, and LNCaP cells.** Cells were fixed and labeled with either the anti-MBOAT4 antibody or with buffer alone, followed by a fluorescently conjugated secondary antibody. In cell expressing hGOAT, immunofluorescence staining is only observed in cells labeled by the primary anti-MBOAT4 antibody. Cells were fixed, stained, and imaged as described in Experimental Procedures.
